# Displayed and Encoded Antigens on Adenovirus Vectors Optimize Humoral and Cellular Immune Responses in Rhesus Macaques

**DOI:** 10.64898/2026.01.29.702655

**Authors:** Zhenyu Li, Matthew D. J. Dicks, Louisa M. Rose, Rebecca A. Russell, Katherine McMahan, Ninaad Lasrado, Joseph Nkolola, Erica Borducchi, Jinyan Liu, Sumi Biswas, Dan H. Barouch

## Abstract

Adenovirus vector-based vaccines were deployed widely during the COVID-19 pandemic. In this study, we explored the potential of displaying an antigen on the surface of the adenovirus capsid as well as encoding an antigen as a transgene in the adenovirus vector to optimize both humoral and cellular immune responses. We show that displaying SARS-CoV-2 Spike receptor biding domain (RBD) on the Ad5 capsid while simultaneously encoding Spike as a transgene induced robust antibody and T cell responses in rhesus macaques. These data demonstrate that adenoviruses can be utilized simultaneously as both nanoparticle scaffolds and viral vectors.

## Main Text

Adenoviruses (Ads) have been widely studied for their potential to be used as vectors to express transgenes of interest for gene therapy and vaccines. During the COVID-19 pandemic, several Ad based vaccines were utilized globally (1–5). Approximately 200 million people received the Johnson & Johnson Ad26.COV2.S vaccine globally, and 2 billion people received the AstraZeneca ChAdOx1 vaccine (4). Ad based vaccines are typically made replication-defective by deleting E1 (5, 6). To increase transgene capacity, E3 can also be deleted (7, 8), and a transgene of interest can be inserted into the viral genome.

Antigen display on nanoparticle scaffolds, using the SpyTag/SpyCatcher protein superglue system (9), has been demonstrated with various vaccine platforms such as hepatitis B virus and mi3 nanoparticles (8, 10, 11). Recent work showed that a related system, Dogtag/Dogcatcher (12), could be used to display immunogens on the capsid surface of adenovirus serotype 5 (Ad5) vectors, effectively utilizing the Ad capsid as the scaffold for a nanoparticle vaccines (13). DogTag (23 amino acids) was genetically inserted into surface-exposed loops in the adenovirus hexon capsid protein to allow covalent attachment of antigens fused to DogCatcher (a 15 kDa protein domain) on virus particles. Attachment by isopeptide bond formation between DogTag and DogCatcher was achieved by co-incubation of antigen and adenovirus components. In mice, Ad5 decorated with the SARS-CoV-2 Spike receptor binding domain (S^RBD^) protein induced over 10-fold higher SARS-CoV-2 neutralization titers compared to an undecorated Ad5 encoding a Spike (S) transgene (12).

Ad-based vaccines expressing transgene encoded antigens have been shown to elicit durable humoral and cellular immune responses in both preclinical and clinical studies (14). Compared with ferritin nanoparticle vaccines, Ad vectors elicited higher CD8^+^ T cell responses but lower antibody responses in non-human primates (NHPs) (15). In this study, we report the immunogenicity in nonhuman primates of Ad5 vectors that both encode SARS-CoV2 Spike (S) as a transgene and display Spike receptor binding domain (S^RBD^) proteins on the capsid surface. This combined vaccine induced more robust humoral and cellular immune responses than Ad5 vectors that only encoded S transgene or only displayed S^RBD^ proteins.

We conducted a head-to-head immunologic comparison of (i) Ad5 that encodes full-length WA1/2020 SARS-CoV2 S as a transgene [Ad5(S)]; (ii) Ad5 that encodes GFP as a negative control transgene and that displays S^RBD^ protein on the capsid surface [Ad5(GFP):S^RBD^]; and (iii) Ad5 that encodes S as a transgene and that displays S^RBD^ protein on the capsid surface [Ad5(S):S^RBD^] (**Figure 1A**). We immunized 18 rhesus macaques (N=6/group) with 5x10^10^ viral particles (vp) of each of these vaccines by the intramuscular route at week 0 and week 4 and measured humoral and cellular immune responses every 2 weeks.

**Figure 1.**
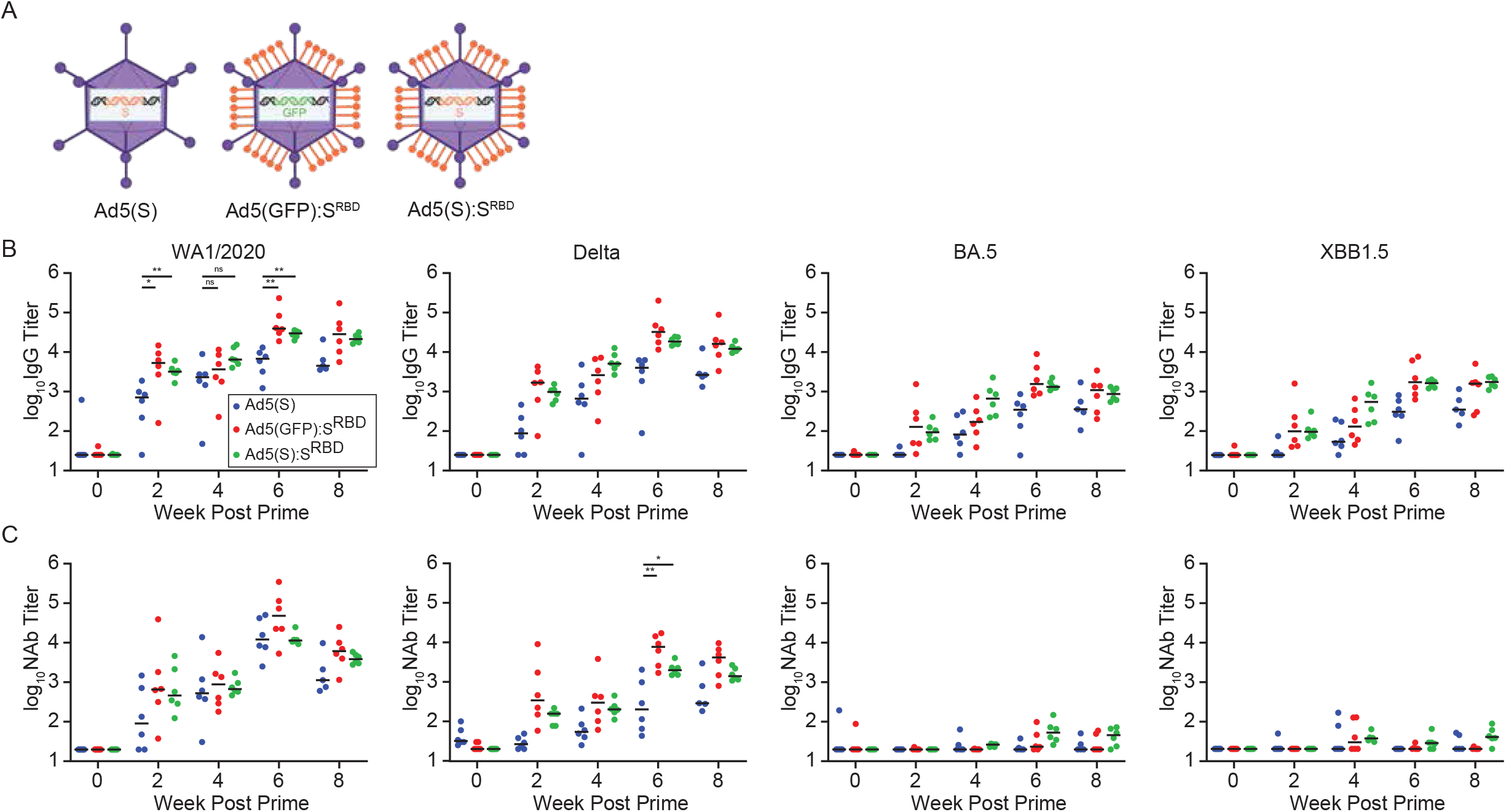
Displaying S^RBD^ on the Ad5 capsid results in higher binding and neutralizing antibody responses. A) Cartoon illustration of vaccines. B) Binding antibody titers against SARS-CoV-2 WA1/2020, Delta, BA.5 and XBB1.5 at weeks 0, 2, 4, 6, 8 (N=18, n=6 per group). Animals were vaccinated at weeks 0 and 4. Median values are shown by the horizontal bars. C) Neutralizing antibody titers against SARS-CoV-2 WA1/2020, Delta, BA.5 and XBB1.5 variants by luciferase-based pseudovirus neutralization assays. Median values are shown by the horizontal bars. Statistical analyses performed by Mann-Whitney U test, *p < 0.05; **p < 0.01; ***p < 0.001; ns, not significant.

All animals developed SARS-CoV-2 WA1/2020, Delta, BA.5, and XBB.1.5 Spike-specific binding antibodies by ELISA by week 2 after the first immunization (**Figure 1B**). By week 6, we observed >10-fold increase in binding antibody titers in all groups. Median binding antibody titers against WA1/2020 Spike at week 6 were 6,789, 39,261 and 29,873 for Ad5(S), Ad5(GFP):S^RBD^ and Ad5(S):S^RBD^ respectively. Consistent with a previous mouse study (12), Ad5(S):S^RBD^ and Ad5(GFP):S^RBD^ elicited 4.4-fold or 5.8-fold higher binding antibody titers than did Ad5(S), respectively, demonstrating the potency of the nanoparticle displayed S^RBD^ antigen for induction of humoral immunity (p=0.0044, p=0.0044 comparing binding antibody titers elicited by Ad5(S):S^RBD^ and Ad5(GFP):S^RBD^ to Ad5(S), respectively, at week 6 for WA1/2020).

Neutralizing antibody (NAb) titers at week 6 against Delta were 202, 7,717 and 1,985 for Ad5(S), Ad5(GFP):S^RBD^ and Ad5(S):S^RBD^ respectively (**Figure 1C**). These data show that Ad5(S):S^RBD^ and Ad5(GFP):S^RBD^ elicited 10-fold or 38-fold higher NAb titers than Ad5(S), respectively (p=0.0152 and p=0.0086, respectively) (**Figure 1C**). As expected, low NAb responses were observed against BA.5 and XBB.1.5 with two immunizations of vaccines encoding WA1/2020 Spike without further boosting, as previously reported (16–18). Additionally, Ad5(GFP):S^RBD^ and Ad5(S):S^RBD^ elicited faster binding antibody kinetics than Ad5(S) from week 2 to week 4 (p=0.0411, p=0.0086, p=0.5887, and p=0.0513 at week 2 and 4 respectively). These results demonstrate that displaying S^RBD^ on the capsid surface results in both higher and faster antibody responses than transgene encoded S only.

We next assessed Spike-specific IFN-γ cellular immune responses by pooled peptide ELISPOT and intracellular cytokine staining (ICS) assays. Ad5(S):S^RBD^ and Ad5(S) elicited higher ELISPOT responses than did Ad5(GFP):S^RBD^. Median WA1/2020 Spike-specific ELISPOT responses were 283, 14 and 589 spot forming cells (SFC) per million PBMC for Ad5(S), Ad5(GFP):S^RBD^ and Ad5(S):S^RBD^ at week 6, respectively, demonstrating the potency of the encoded S transgene for cellular immunity (p=0.0044 and p=0.0044 comparing WA1/2020 Spike-specific ELISPOT responses elicited by Ad5(S):S^RBD^ and Ad5(S) to Ad5(GFP):S^RBD^, respectively) (**Figure 2A**) (12). Similar results were observed for both CD8^+^ and CD4^+^ T cell responses by ICS assays (**Figure 2B, C**). Moreover, T cell responses were highly cross-reactive against multiple SARS-CoV-2 variants, consistent with previous reports (17, 19). These data showe that the S transgene resulted in higher cellular immune responses than capsid displayed S^RBD^.

**Figure 2.**
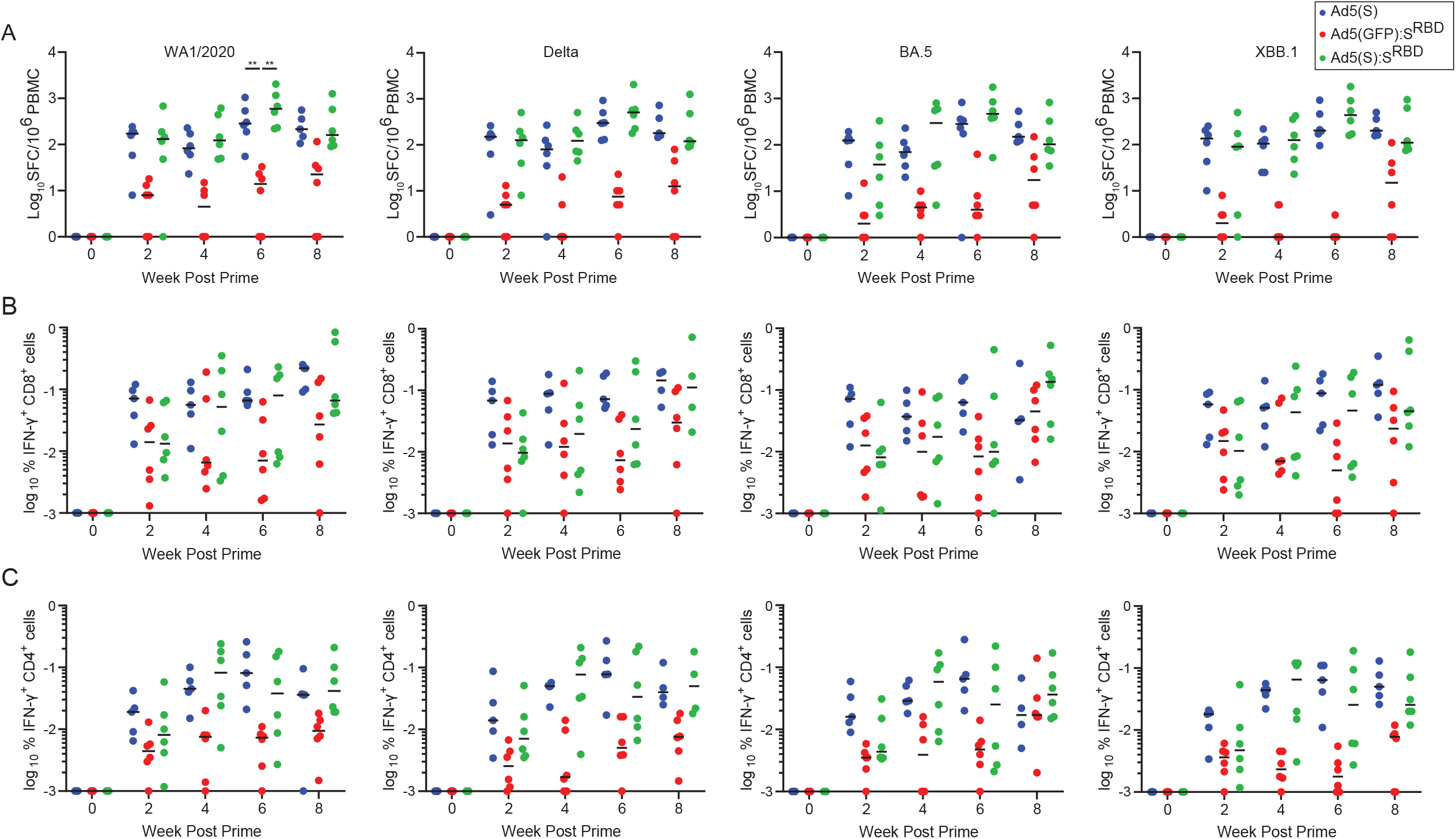
Encoding S as a transgene results in higher T cell response. A) Spike-specific pooled peptide IFN-γ ELISPOT responses to SARS-CoV-2 WA1/2020, Delta, BA.5 and XBB1.5 at weeks 0, 2, 4, 6, 8 (N=18, n=6 per group). Animals were vaccinated at weeks 0 and 4. Median values are shown by the horizontal bars. Spike-specific pooled peptide IFN-γ CD8^+^ T (B) cells and CD4^+^ T cells (C) were assessed by intracellular cytokine staining. Statistical analyses performed by Mann-Whitney U test, *p < 0.05; **p < 0.01; ***p < 0.001; ns, not significant.

Taken together, we show in nonhuman primates that display of S^RBD^ on an adenovirus capsid drives higher antibody responses and that transgene encoded S induces higher T cell responses. The combined vaccine Ad5(S):S^RBD^ that both displays S^RBD^ on the capsid surface and that encodes S as a transgene induced both robust humoral and robust cellular immune responses. These data demonstrate that Ad5 can function simultaneously as a nanoparticle scaffold for a protein immunogen and as a classical viral vector for an encoded transgene. Future studies could evaluate the potential of displaying B cell immunogens (e.g. influenza hemagglutinin) on the capsid surface and encoding T cell immunogens (e.g. influenza matrix or nucleoprotein) as a transgene. Adenovirus capsid decoration and transgene delivery have the potential to be a highly versatile and customizable platform, with applications in seasonal vaccination, pandemic preparedness, and personalized medicine.

### Animals

18 outbred male and female rhesus macaques was housed at Alphagenesis Inc. Animals were randomly allocated and the study was conducted in compliance with all relevant local, state and federal regulations, and all animal procedures were approved by the Alphageneis Institutional Animal Care and Use Committee. Each animal was immunized with 5x10^10^ viral particles (vp) of each vaccine at week 0 and week 4. Blood samples were collected every 2 weeks for the first two months of study and every 4 weeks for the remainder of the study.

### Vaccines

Construction of vaccines has been described previously (12). Briefly, WA1/2020 Spike or green fluorescent protein (GFP) was inserted into E1/E3-deleted Ad5 after a cytomegalovirus immediate-early promoter containing a tetracycline operator sequence as previously described. DogTag (DIPATYEFTDGKHYITNEPIPPK) flanked by GSGGSG linkers was inserted into the hexon HVR5 loop through BAC GalK recombineering (12). Viruses were rescued and amplified in E1-complementing human embryonic kidney (HEK) 293 cell lines, either HEK293A (GFP expressing Ad) or HEK293TREX (Invitrogen) (S expressing Ad). Viruses were purified via CsCl gradient ultracentrifugation. Purified virus was stored in sucrose storage buffer (10mM Tris-HCl, 7.5% w/v sucrose, pH7.8) at -80°C. SARS-CoV-2 Spike receptor binding domain (RBD) fused to dogcatcher (Dogcatcher-S^RBD^) (WA1/2020, residues 319-532) was cloned into a mammalian protein expression plasmid, produced in ExpiCHO-S cells and purified via C-tag affinity chromatography (Thermo Fisher) as previously described (12). Purified protein were dialyzed into tris-buffered saline (TBS). DogTag/DogCatcher conjugation to attach DogCatcher-S^RBD^ to adenovirus particles was performed as previously described (12). Excess ligand was removed by SpectraPor dialysis cassettes (300-kDa MWCO) and decorated particles were stored at -80°C (12). All vaccines were filter sterilized (0.22µm, Millex-GP) prior to storage.

### ELISA

SARS-CoV-2 spike specific binding antibodies in serum were assessed by ELISA. 96-well plates were coated with 1 μg/mL of similarly produced SARS-CoV-2 WA1/2020, Delta, BA.5 or XBB1.5 Spike protein receptor binding domain in 1× Dulbecco phosphate-buffered saline (DPBS) and incubated at 4 °C overnight. After incubation, plates were washed once with wash buffer (0.05% Tween 20 in 1× DPBS) and blocked with 350 μL of casein block solution per well for 2 to 3 hours at room temperature. Following incubation, block solution was removed and plates were blotted dry. Heat-inactivated serum was serially diluted in Casein block buffer containing wells. Plates were incubated at room temperature for 1 hour, prior to 3 more washes and a 1-hour incubation with a 1:1000 dilution of anti–macaque IgG horseradish peroxidase (HRP) (Nonhuman Primate Reagent Resource) at room temperature in the dark. Plates were washed 3 more times, and 100 μL of SeraCare KPL TMB SureBlue Start solution was added to each well; plate development was halted by adding 100 μL of SeraCare KPL TMB Stop solution per well. The absorbance at 450 nm, with a reference at 650nm, was recorded with a VersaMax microplate reader (Molecular Devices). For each sample, the ELISA end point titer was calculated using a 4-parameter logistic curve fit to calculate the reciprocal serum dilution that yields an absorbance value of 0.2. Interpolated end point titers were reported.

### Pseudovirus Neutralization Assay

Heat-inactivated sera were tested for neutralization against WA1/2020, Delta, BA.5 and XBB1.5 SARS-CoV2 variants as previously described using lentivirus-based pseudotyped virus encoding a luciferase reporter gene. Briefly, human embryonic kidney HEK293T cells (ATCC CRL_3216) were co-transfected with a luciferase reporter plasmid (pLenti-CMV Puro-Luc, Addgene), packaging construct psPAX2 (AIDS Resource and Reagent Program) and Spike protein expressing pcDNA3.1-SARS-CoV-2 SΔCT using lipofectamine 2000 (Thermo Fisher). The pseudovirus containing supernatants were collected two days after transfection and filtered through a 0.45-μm Filter. To determine neutralization titers, 3-fold dilutions of the sera were pre-incubated with the same volume of pseudovirus for 1h at 37°C before the sera/pseudovirus-mix was transferred to HEK293T-ACE2 cells seeded the day before (2 × 10^4^ cells/well). Approximately 48h later, cells were lysed using Steady-Glo Luciferase System (Promega) according to manufacturer’s instructions. Neutralization titer was determined as the sera dilution in which 50% of relative light units were observed, when 100% relative light units were set to be virus only and 0% were set to be with neither virus nor sera.

### Enzyme Linked ImmunoSPOT Assay (ELISPOT)

Peptide pools were 15 amino acid peptides overlapping by 11 amino acids spanning the SARS-CoV-2 WA1/2020, Delta, BA.5 or XBB1.5 spike proteins (21st Century Biochemicals). IFN-γ ELISPOT was performed on PBMCs. ELISPOT plates were coated with mouse anti-human IFN-γ monoclonal antibody from BD Pharmingen at 0.5 µg per well and incubated overnight at 4 °C. Plates were washed with DPBS containing 0.25% Tween 20 and blocked with R10 media (500 mL RPM1460 with 55 mL PFBS and 5.5 mL of 100X penicillin-streptomycin) for 1 to 4 hours at 37°C. SARS-CoV-2 pooled S peptides (1 µg/mL) and cells (2×105/well) were added to the plate and incubated for 18 to 24 hours at 37°C. All steps following this incubation were performed at room temperature. The plates were washed with coulter buffer and incubated for 2 hours with biotinylated anti-human IFN-γ antibody from U-CyTech Biosciences (1 µg/mL in each well). The plates were washed a second time and incubated for 2 hours with streptavidin-AP antibody from Southern Biotechnology (2 µg/mL in each well). The final wash was followed by the addition of warmed and filtered nitro blue tetrazolium chloride/5-bromo-4-chloro 3 ‘indolyl phosphate p-toluidine salt (BCIP/NBT chromogen) substrate solution from Pierce. The chromogen was discarded, and the plates were washed with water and dried out of direct light for 24 hours. Plates were scanned and counted on a KS Elispot Reader. Results are expressed with a background subtraction.

### Intracellular Cytokine Staining

CD4+ and CD8+ T cell responses were quantitated by pooled peptide-stimulated intracellular cytokine staining (ICS) assays. Peptide pools were the same as mentioned above in EliSpot assay. 10^6^ peripheral blood mononuclear cells well were re-suspended in 100 μL of R10 media supplemented with CD49d monoclonal antibody (1 μg/mL) and CD28 monoclonal antibody (1 μg/mL). Each sample was assessed with mock (100 μL of R10 plus 0.5% DMSO; background control), peptides (2 μg/mL), and/or 10 pg/mL phorbol myristate acetate (PMA) and 1 μg/mL ionomycin (Sigma-Aldrich) (100μL; positive control) and incubated at 37°C for 1 h. After incubation, 0.25 μL of GolgiStop and 0.25 μL of GolgiPlug in 50 μL of R10 was added to each well and incubated at 37°C for 8 h and then held at 4°C overnight. The next day, the cells were washed twice with DPBS, stained with aqua live/dead dye for 10 mins and then stained with predetermined titers of monoclonal antibodies against CD279 (clone EH12.1, BB700), CD38 (clone OKT10, PE), CD28 (clone 28.2, PE CY5), CD4 (clone L200, BV510), CD95 (clone DX2, BUV737), CD8 (clone SK1, BUV805) for 30 min. Cells were then washed twice with 2% FBS/DPBS buffer and incubated for 15 min with 200 μL of BD CytoFix/CytoPerm Fixation/Permeabilization solution. Cells were washed twice with 1X Perm Wash buffer (BD Perm/WashTM Buffer 10X in the CytoFix/CytoPerm Fixation/ Permeabilization kit diluted with MilliQ water and passed through 0.22μm filter) and stained with intracellularly with monoclonal antibodies against Ki67 (clone B56, FITC), CD69 (clone TP1.55.3, ECD), IL10 (clone JES3-9D7, PE CY7), IL13 (clone JES10-5A2, BV421), TNF-α (clone Mab11, BV650), IL4 (clone MP4-25D2, BV711), IFN-γ (clone B27; BUV395), CD45 (clone D058-1283, BUV615), IL2 (clone MQ1-17H12, APC), CD3 (clone SP34.2, Alexa 700) for 30 min. Cells were washed twice with 1X Perm Wash buffer and fixed with 250μL of freshly prepared 1.5% formaldehyde. Fixed cells were transferred to 96-well round bottom plate and analyzed by BD FACSymphony™ system. Data were analyzed using FlowJo v9.9.

## Statistical Analysis

Data were plotted and analyzed by GraphPad Prism v10.6.1. All statistical analysis were compared using two-sided Mann-Whitney U-tests with Holm-Bonferroni correction. P<0.05 was considered to be significant.

## Acknowledgements

We thank Darren Ty, Kristin Gotthardt, Nicole Hachman and Jessica Miller for their help in intracellular staining, EliSpot and pseudovirus neutralization assays. We acknowledge SpyBiotech for funding.

## Conflicts of Interest

M.D.J.D, L.M.R., R.A.R. and S.B. are employees of Spybiotech. The other authors declare no conflicts of interest.

